# Prenatal Alcohol Exposure Disrupts γ-Secretase Activity and Impairs Learning and Memory in Wild-Type and 3xTg-AD Mice

**DOI:** 10.64898/2026.06.11.731622

**Authors:** Paola Montenegro, Rachel Kim, Mikayla Zedek, Marianna Chicas, Pamela Yeh, Hermes Yeh

## Abstract

Although prenatal alcohol exposure (PAE) has been proposed as an early-life risk factor for Alzheimer’s disease and related dementias (AD/ADRD), the mechanistic underpinnings are underexplored. Mutations in the *Presenilin* genes contribute to AD/ADRD, with Presenilin 1 acting as the catalytic subunit of the γ-secretase complex responsible for cleaving Notch and amyloid precursor protein (APP). We hypothesized that PAE disrupts γ-secretase activity during brain development, which persists and is associated with behavioral deficits later in life. Pregnant wild-type B6129 and 3xTg-AD mice were fed an ethanol-containing liquid diet during gestational days 13–15. From birth to adulthood, PAE increased APP C-terminal fragments and Notch intracellular domain (NICD) levels in cortical lysates. These changes were accompanied by impaired hippocampal-dependent learning and memory in wild-type mice at 3 and 6 months of age and exacerbated behavioral deficits in 4-month-old 3xTg-AD mice. Our findings provide the first mechanistic insight linking PAE to AD/ADRD vulnerability.

## Introduction

There has been great interest in unraveling environmental risk factors that contribute to advancing our appreciation of the pathoetiology and treatment of Alzheimer’s disease and Alzheimer’s disease–related dementias (AD/ADRD) (Borenstein et al., 2006). To date, while such studies have largely focused on risk factors that are prevalent in adulthood, they also underscore that potential early-life risk factors have been unexplored (Yu et al., 2020).

Why consider early-life events as risk factors when the lore is that AD is an adult-onset neurodegenerative disease? Neurodevelopment and aging are stages in the continuum of life; neurodevelopment begins in the embryo, continues through childhood, adolescence, and tapers off in young adulthood. Disruption or impairment of neurodevelopmental processes at any point in this continuum can lead to indelible lifelong disabilities, many of which are reminiscent of brain abnormalities or impaired cognition seen in neurodegenerative disorders. Along this line of thought, neurodevelopmental disorders such as autism spectrum have been reported to be associated with AD, a neurodegenerative disorder (Nadeem et al., 2021).

Early-life environmental insults can leave molecular and cellular imprints that persist long after development (Basha et al., 2005). The developing brain is particularly susceptible to environmental insults that enhance vulnerability to neurodegenerative diseases throughout the lifespan (Sarkar et al., 2019; Tartaglione et al., 2016). Prenatal alcohol exposure (PAE) may be one such insult, postulated in preclinical studies to be an early-life risk factor that contributes to increased vulnerability to developing AD/ADRD later in life (Borenstein et al., 2006; Huang et al., 2023; Lahiri et al., 2008; Tousley et al., 2023). Indeed, PAE is recognized clinically as a potential early-life contributor to the development of neurobehavioral disorders (Coles et al., 2020; Hagan et al., 2016). Importantly, binge drinking is a common mode of PAE, particularly during the first trimester of gestation, before pregnancy is recognized, but just when neurogenesis and brain patterning peak (Dejong et al., 2019; Denny et al., 2019; Tough et al., 2006). Binge-like maternal alcohol consumption produces more severe neurodevelopmental outcomes than chronic low-level exposure, highlighting the importance of exposure pattern in determining long-term risk (Dejong et al., 2019; Maier & West, 2001; Muggli et al., 2024).

Prenatal alcohol exposure adversely affects key neurodevelopmental processes, including neurogenesis, neuronal migration, and synaptogenesis (Coles et al., 2020). These disruptions alter the formation and maturation of neural circuits, leading to persistent structural and functional abnormalities. Clinically, such deficits manifest themselves as long-term cognitive impairments, including deficits in learning, memory, and executive function, which are core features observed in individuals with Fetal Alcohol Spectrum Disorders (FASD) (Franklin et al., 2008; Mattson et al., 2019; Rasmussen, 2005). Although the behavioral and cognitive consequences of PAE are well documented (Chung et al., 2021), the molecular mechanisms linking early binge-like alcohol exposure to later-life neurodegenerative processes remain poorly understood. The present study is grounded in the perspective that indelible neurodevelopmental alterations may increase susceptibility to the development of neurodegenerative disorders, including AD/ADRD.

AD/ADRD are the most prevalent neurodegenerative disorders, characterized by progressive cognitive decline, synaptic dysfunction, and neuronal loss (Sheppard & Coleman, 2020). Mutations in the *Presenilin* genes (*PSEN1* and *PSEN2*) are among the most common causes of familial AD (Kelleher & Shen, 2017). Presenilin-1 (PS1) functions as the catalytic subunit of the γ-secretase complex, a protease that cleaves type I transmembrane proteins, including the amyloid precursor protein (APP) and Notch (Schroeter et al., 2003). For γ-secretase to become catalytically active, PS1 undergoes an endoproteolytic self-cleavage that generates stable N-terminal (NTF) and C-terminal (CTF) fragments (Gertsik et al., 2015; Schroeter et al., 2003; Wolfe, 2019). The γ-secretase complex, through its proteolytic activity, regulates key signaling pathways involved in neurogenesis, neuronal differentiation, and synaptic plasticity during brain development, and supports neuronal survival and synaptic function in adulthood (Jurisch-Yaksi et al., 2013; Wolfe, 2019). Disruptions in γ-secretase function during early development, driven by genetic mutations in *PSEN1*, result in loss-of-function alterations that impair Notch signaling and APP processing, leading to defective neurogenesis, altered neuronal differentiation, and long-lasting synaptic deficits (Handler et al., 2000; Zhou et al., 2011). PAE appears to converge on similar neurodevelopmental outcomes. PAE has been shown to interfere with key developmental signaling pathways, including those regulated by γ-secretase substrates, resulting in reduced neuronal proliferation, impaired neuronal migration and circuit formation, and persistent deficits in synaptic plasticity (Delatour et al., 2019; Guerri, 2002; Miller, 1993, 1996; Sarmah et al., 2016; Tong et al., 2013). These converging effects suggest that PAE may phenocopy aspects of γ-secretase dysfunction during brain development, thereby establishing a potential mechanistic framework through which early-life environmental exposures can increase vulnerability to AD/ADRD later in life.

In the context of APP processing, β-secretase (BACE1) cleavage first generates the membrane-bound APP C-terminal fragment (C99), which is subsequently processed by γ-secretase to produce amyloid-β (Aβ) peptides and the APP intracellular domain (Hur, 2022). Disruption of γ-secretase activity interferes with this process and leads to the intracellular accumulation of APP C-terminal fragments (APP CTFs), including C99. Elevated levels of APP CTFs are widely used as a biochemical indicator of reduced γ-secretase activity (Heilig et al., 2010; Montenegro et al., 2023; H. Yu et al., 2001). Importantly, the accumulation of C99 itself is increasingly recognized as a pathogenic contributor to AD, as it promotes endosomal dysfunction, synaptic deficits, and neurotoxicity independent of extracellular Aβ deposition (Bourgeois et al., 2018; Pulina et al., 2020). Supporting this, experimental studies in adult murine models have shown that ethanol exposure enhances amyloidogenic APP processing by increasing BACE1 activity, leading to elevated production of C99 and Aβ peptides (Huang et al., 2018). Ethanol-induced oxidative stress and membrane alterations have been proposed to shift APP processing toward the amyloidogenic pathway, thereby promoting the accumulation of neurotoxic APP-derived fragments (Huang et al., 2018). Consistent with these findings, studies have shown that excessive alcohol consumption disrupts the processing of APP, Aβ, and the expression of enzymes involved in these processes (Gong et al., 2021; Kim et al., 2011; Renau-Piqueras et al., 1997; Venkataraman et al., 2017). Moreover, multiple studies have linked PAE to AD-like pathology in mouse models (Alhowail, 2022; Canales et al., 2013; Gerlikhman et al., 2023; Tousley et al., 2023), suggesting that APP processing, hence γ-secretase activity, may be disrupted during neurodevelopment.

In this study, we hypothesize that PAE alters γ-secretase activity early during brain development, resulting in disrupted APP and Notch processing and increased vulnerability to learning and memory deficits in adulthood. To this end, we employed a binge-type PAE paradigm in both wild-type B6129 and 3xTg-AD mice (Fig. 1A), the latter harboring three human AD/ADRD-associated mutations: *PSEN1* M146V, *Tau P301L,* and *APP K670_M671delinsNL* (Oddo et al., 2003). These two mouse models served complementary purposes in this study. Wild-type B6129 mice were used to characterize the effects of PAE on γ-secretase activity and to investigate the long-term consequences of PAE on learning and memory. The 3xTg-AD model served both as a pathological reference for contextualizing PAE-induced changes in wild-type mice relative to an established genetic model of AD and as an experimental model to directly investigate how PAE influences AD-related mutations. This comparative approach allowed us to examine whether PAE-induced alterations overlap with or differ from those associated with AD-related mutations, while also exploring the prospect that PAE may modify the progression or severity of AD-related phenotypes in a genetically susceptible background. We conducted biochemical and histological analyses to assess the effects of PAE on γ-secretase function, as well as behavioral analyses to assess learning and memory. Our findings provide novel insights into how early-life alcohol exposure contributes to AD/ADRD pathogenesis, highlighting γ-secretase dysfunction as a potential mechanistic link between PAE and memory decline throughout life span.

**Figure 1.**
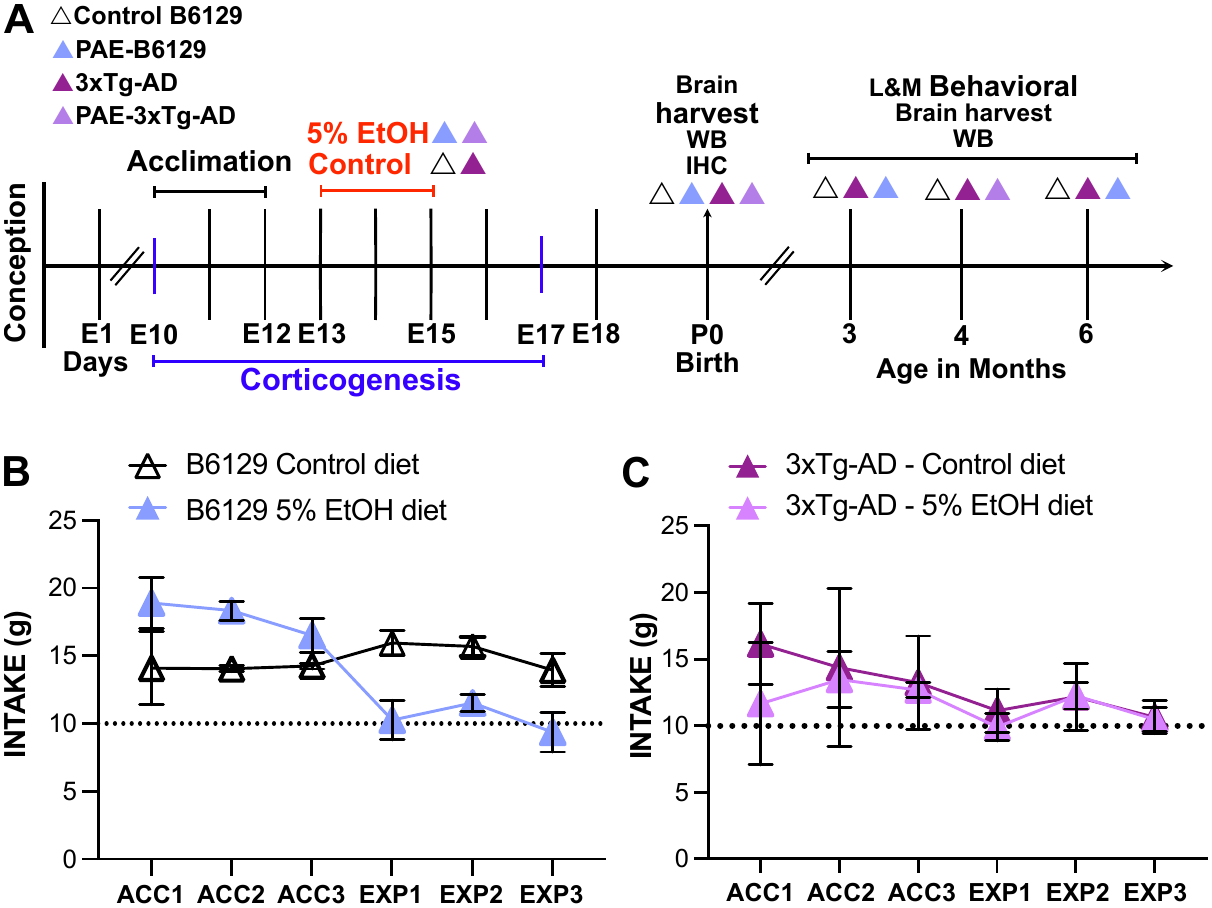
Experimental timeline and dams’ average alcohol intake. **(A)** The timeline illustrates the experimental design, including an acclimation period to a liquid diet from embryonic days (E) 10 to E12, followed by prenatal alcohol exposure from E13 to E15, at the peak of corticogenesis. Biochemical, histological, and behavioral assessments were performed at different postnatal ages. At least four dams per experimental group (color-coded in the top legend) received either a 5% ethanol diet or an isocaloric control diet. Within each litter, approximately half of the offspring underwent brain harvesting at birth (P0), while the remaining offspring were maintained until the indicated ages (in months). Sample sizes for each experimental group are reported in the corresponding results figure. **(B-C)** Liquid diet consumption by B6129 dams (B) and 3xTg-AD dams (C) during acclimation (ACC) and experimental (EXP) periods. Both groups of dams consumed more than 10 g of either the isocaloric control diet or the 5% ethanol (EtOH) diet, indicating that both groups were exposed to an equivalent ethanol concentration.

## Results

### PAE impairs γ-secretase activity in neonatal wild-type B6129 and 3xTg-AD mice

Amyloid precursor protein C-terminal fragments (APP CTFs) and the Notch intracellular domain (NICD) are key readouts of γ-secretase activity, with APP representing the immediate substrate for γ-secretase cleavage and NICD a direct product of γ-secretase–mediated processing of the Notch receptor (Schroeter et al., 2003). In this light, we asked whether prenatal alcohol exposure (PAE) alters γ-secretase–dependent processing during cortical development by quantifying levels of APP CTFs and NICD in lysates harvested from the cortex towards the end stage of embryonic corticogenesis at postnatal day 0 (P0, birth).

Figure 2 illustrates Western blot data assessing γ-secretase–related processing of PS1 and APP expression in cortex lysates derived from P0 wild-type B6129 and 3xTg-AD mice with and without PAE. The summary graphs in Figure 2C and 2F indicate disrupted APP processing in wild-type B6129 and 3xTg-AD mice, respectively, exposed prenatally to ethanol (PAE-B6129 and PAE-3xTg-AD) relative to their isocaloric liquid diet controls. In wild-type B6129 mice, PAE significantly increased levels of APP CTFs up to 31.21% in the cortex relative to their non-PAE control cohorts (p = 0.0026, unpaired Student’s t-test, Fig. 2A and 2C). NICD levels were also significantly elevated up to 34.14% in PAE-B6129 mice relative to their non-PAE controls (p = 0.0315, unpaired Student’s t-test, Fig. 2B and 2C). In P0 3xTg-AD mice, cortical APP CTFs levels were significantly increased in both non-PAE (36.20%, p = 0.0395, one-way ANOVA with Tukey’s post hoc multiple comparisons) and PAE (64.35%, p = 0.0011, one-way ANOVA with Tukey’s post hoc multiple comparisons) groups compared to non-PAE B6129 controls (Fig. 2D and 2F). Similarly, NICD levels were significantly elevated in PAE-3xTg-AD mouse cortex (53.96%, p=0.0087), while non-PAE 3xTg-AD showed a trend towards the accumulation of NICD; however, this difference did not reach statistical significance (p = 0.07, one-way ANOVA with Tukey’s post hoc multiple comparisons, Figs. 2E and 2F), compared to non-PAE B6129 controls. The increase in both cortical APP CTFs and NICD levels suggests that PAE affects the normal regulation of the γ-secretase proteolytic pathway at birth, resulting in increased intracellular accumulation. Importantly, the supranormal levels of APP CTFs and NICD observed in PAE-3xTg-AD mice suggest that PAE exacerbates the dysregulation of APP and Notch processing beyond that driven by AD-linked mutations.

**Figure 2.**
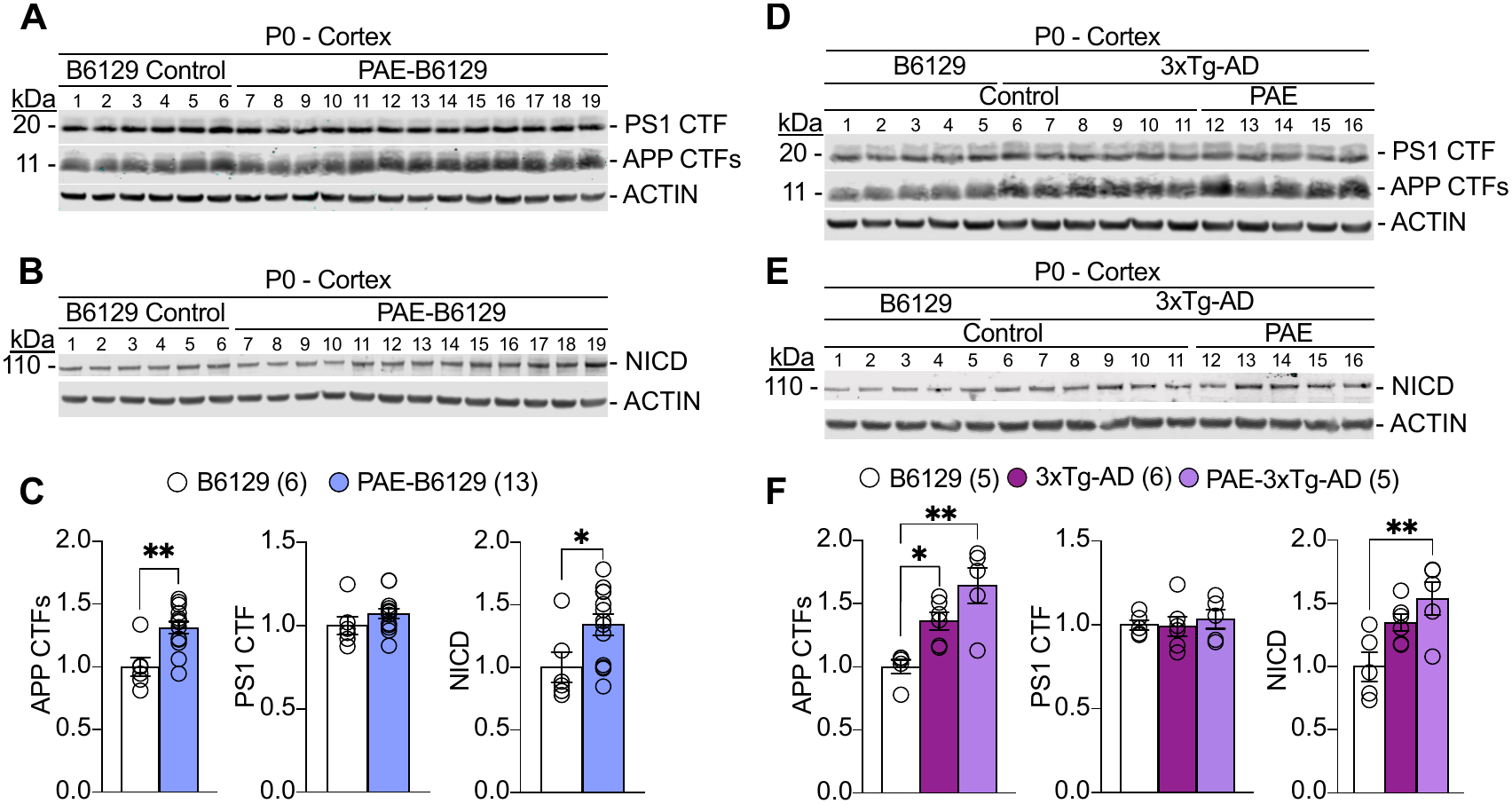
PAE reduces γ-secretase activity in B6129 and 3xTg-AD mice at birth relative to non-exposed wild-type mice. **(A)** Western analysis of cortical lysates from B6129 controls (lanes 1-6) and PAE-B6129 (lanes 7-19) mice at postnatal day 0 (P0) shows no change in PS1 CTF levels between groups, and increased levels of APP CTFs in PAE-B6129 mice compared to B6129 controls. **(B)** Western blot analysis of cortical lysates from B6129 controls (lanes 1-6) and PAE-B6129 (lanes 7-19) mice at P0 shows increased levels of NICD in PAE-B6129 mice compared to controls. **(C)** Quantification of APP CTFs, PS1 CTF, and NICD shows that levels of APP CTFs (p = 0.0026) and NICD (p = 0.0315) are significantly increased compared to non-alcohol-exposed B6129. Quantification of PS1 CTF levels shows no significant differences between PAE-B6129 and B6129 controls (p = 0.2269). Unpaired Student’s t-test. **(D)** Western analysis of cortical lysates from B6129 controls (lanes 1-5), 3xTg-AD controls (lanes 6-11), and PAE-3xTg-AD (lanes 12-16) mice at P0 shows no change in PS1 CTF levels between groups, and increased levels of APP CTFs in 3xTg-AD controls and PAE-3xTg-AD mice compared to B6129 controls. **(E)** Western analysis of cortical lysates from B6129 controls (lanes 1-5), 3xTg-AD controls (lanes 6-11), and PAE-3xTg-AD (lanes 12-16) mice at P0 shows increased levels of NICD in 3xTg-AD controls and PAE-3xTg-AD PAE-B6129 mice compared to B6129 controls. **(F)** Quantification of APP CTFs, PS1 CTF, and NICD shows that levels of APP CTFs in 3xTg-AD controls (p = 0.0395) and in PAE-3xTg-AD (p = 0.0011) were significantly higher compared to B6129 controls. Quantification of PS1 CTF levels shows no significant differences between 3xTg-AD controls (p = 0.9923), PAE-3xTg-AD (p = 0.8980), and B6129 controls. Levels of NICD in PAE-3xTg-AD (p = 0.0087) are significantly higher compared to B6129 controls. One-way ANOVA with Tukey’s post hoc multiple comparisons. All data represent mean ± SEM. *p < 0.05, **p < 0.01. Sample size is indicated in parentheses next to each experimental group label.

We then asked whether the observed alterations in γ-secretase activity are associated with changes in the γ-secretase complex itself by examining the expression of the Presenilin 1 C-terminal fragment (PS1 CTF), a key functional component of the γ-secretase complex (Fig. 2A and 2D). As summarized in Figures 2C and 2F, PS1 CTF levels were unchanged in PAE-B6129 (p = 0.2269, unpaired Student’s t-test), 3xTg-AD (p = 0.9923, one-way ANOVA with Tukey’s post hoc multiple comparisons), and PAE-3xTg-AD (p = 0.8980, one-way ANOVA with Tukey’s post hoc multiple comparisons) compared to wild-type controls, suggesting that PAE-induced alterations in γ-secretase activity are unlikely to result from changes in the abundance of its catalytic subunit.

To gain insight into how PAE influences the spatial distribution of the APP CTFs protein in brain regions that are vulnerable in patients with AD, we assessed immunohistochemically the intracellular accumulation of the APP CTFs protein in cortical and hippocampal sagittal sections derived from P0 wild-type B6129, PAE-B6129, 3xTg-AD, and PAE-3xTg-AD mice. APP CTFs were widely distributed throughout the cortical layers (Fig. 3, top panels). Quantification of mean fluorescence intensity revealed that the strongest immunoreactivity was observed across layers II–VI, with significant increases detected in the PAE-B6129 up to 32.54% (p = 0.0216, one-way ANOVA with Tukey’s post hoc multiple comparisons) and PAE-3xTg-AD up to 34.01% (p = 0.0261, one-way ANOVA with Tukey’s post hoc multiple comparisons) mice. In the hippocampus, APP CTFs were prominently localized within the CA regions, notably in the CA1 and CA2/3 subregions, compared to age-matched wild-type B6129 controls (Fig. 3, bottom panels; PAE-B6129, 28.50%, p = 0.0030; 3xTg-AD, 26.05%, p = 0.0088; and PAE-3xTg-AD, 26.46%, p = 0.0105. One-way ANOVA with Tukey’s post hoc multiple comparisons.). Thus, brain regions essential for learning, memory, and the formation of cortical networks are already affected by altered APP processing by birth in the cortex and hippocampus. Together, these findings demonstrate that PAE is sufficient to induce early dysregulation of γ-secretase–related pathways in a wild-type B6129 background, as reflected by increases in APP CTFs and NICD at birth. In the presence of AD-associated mutations, this disruption is exacerbated, leading to greater accumulation of APP processing intermediaries. Thus, prenatal alcohol exposure may act as an early-life modifier of molecular pathways linked to Alzheimer’s disease and related dementias, including impaired learning and memory behavior.

**Figure 3.**
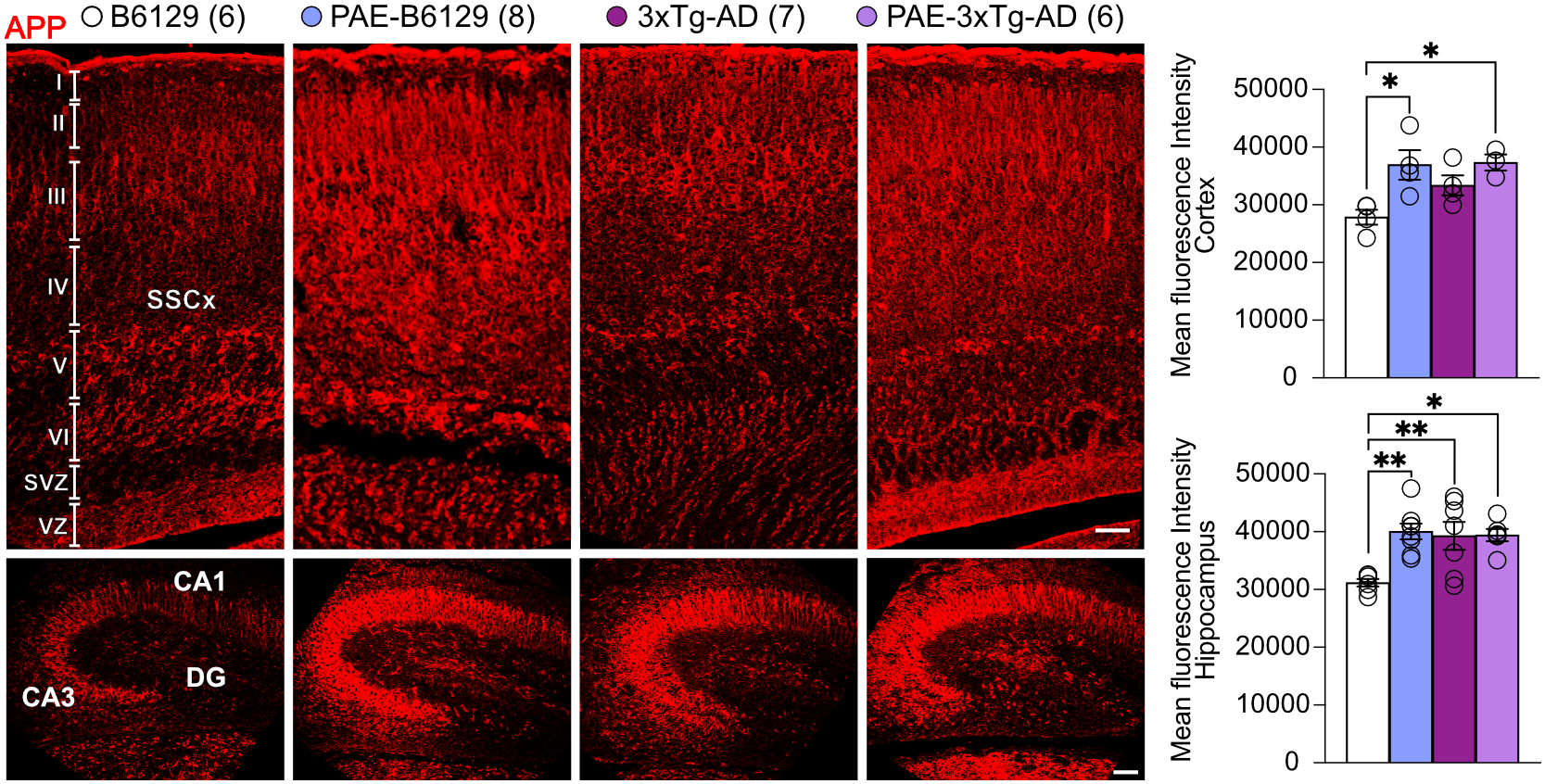
PAE-B6129 and PAE-3xTg-AD mice show high expression of APP CTFs in cortical regions at birth. (Top Panel) Representative images of cortical sagittal sections immunoassayed using antibodies specific for the C-terminal fragment of APP. Accumulation of APP CTFs is evident in the somatosensory cortex (SSCx), in layers II to VI, of PAE-B6129, 3xTg-AD, and PAE-3xTg-AD mice compared to B6129 controls. Quantification of fluorescence signal shows a significant increase in PAE-B6129 (p = 0.0216) and PAE-3xTg-AD (p = 0.0261) compared to B6129 controls. **(Bottom Panel)** Hippocampal sagittal representative images showing accumulation of APP CTFs mostly in CA1, CA2/3, and modest increased immunoreactivity in dentate gyrus (DG). Quantification of fluorescence signal shows a significant increase in PAE-B6129 (p = 0.0030), 3xTg-AD (p = 0.0088), and PAE-3xTg-AD (p = 0.0105) compared to B6129 controls. One-way ANOVA with Tukey’s post hoc multiple comparisons. All data represent mean ± SEM. *p < 0.05, **p < 0.01. Sample size indicated in parentheses next to each experimental group label. Scale bars = 50 μm.

### PAE impairs learning and memory and γ-secretase-related activity in adult wild-type B6129 mice

To determine if the disruption of γ-secretase–mediated processing observed at birth persists and is associated with learning and memory impairments in adulthood, hippocampal-dependent spatial learning was first assessed using the Morris water maze in wild-type B6129 mice at 3 and 6 months of age with or without PAE. The levels of APP CTFs and PS1 CTFs were analyzed using Western blot in cortical and hippocampal lysates derived from the same behaviorally tested mice.

Figure 4 illustrates the timeline of the Morris water maze regime (Fig. 4A) and behavioral data (Fig. 4B) derived from 3-month-old young adult B6129 mice with and without PAE. During hidden platform training in the Morris water maze, PAE-B6129 mice showed a significant longer escape latencies compared to control B6129 mice (p = 0.0036, two-way ANOVA with Tukey’s post hoc multiple comparisons of the main columns means), indicating impaired spatial learning in the acquisition phase (Fig. 4A). These deficits in B6129 mice with PAE were detected at by 3 months of age when 3xTg-AD mice typically are presymptomatic for AD pathology, suggesting that PAE alone is sufficient to induce impaired hippocampal-dependent learning.

**Figure 4.**
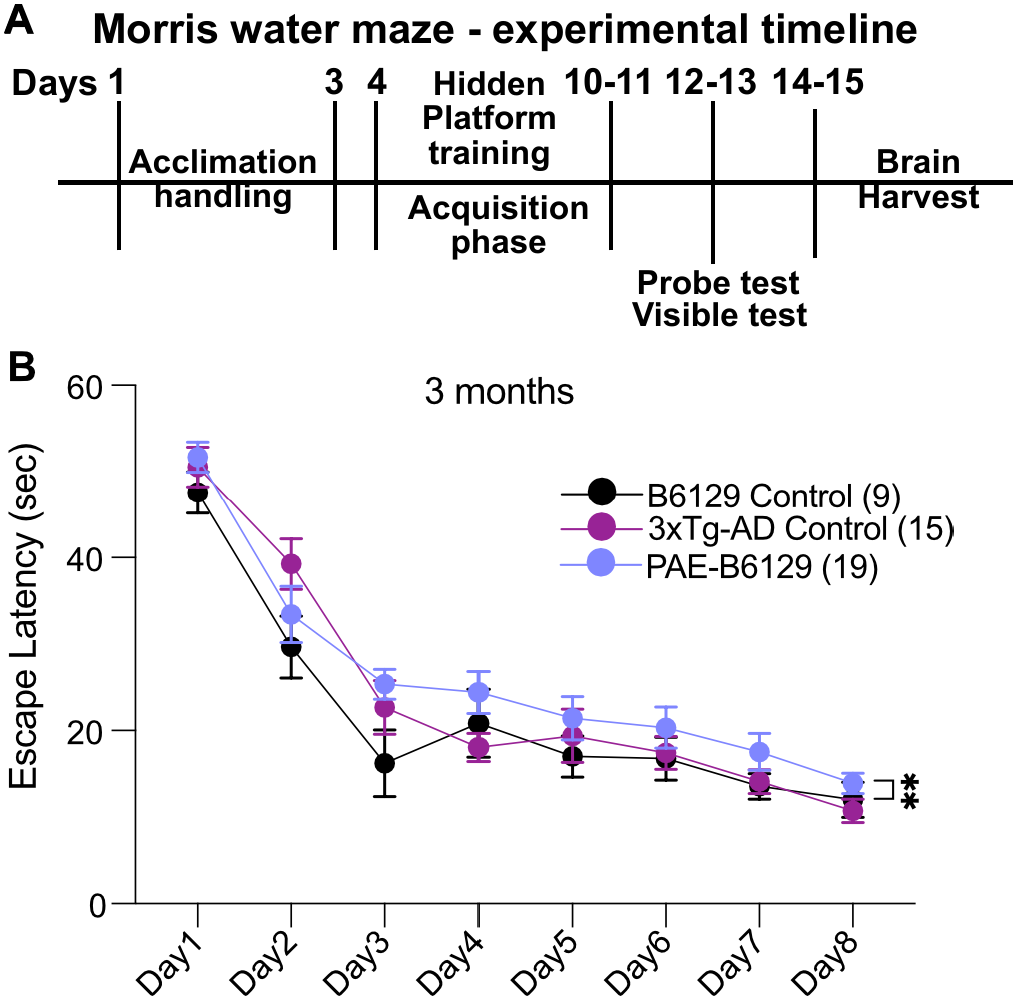
PAE compromises learning and memory in B6129 mice at 3 months of age. **(A)** Morris water maze experimental timeline: Mice are individually acclimated to handling during days 1 to 3. The hidden platform training and acquisition phase is conducted from days 4 to 10-11, for a total of 7-8 days. Probe and visible platform tests are performed one day after completion of the acquisition phase, on days 12-13. Brains are harvested immediately following the behavioral study, on days 14-15. **(B)** During the hidden platform version of the Morris water maze test, PAE-B6129 mice at 3 months of age display significantly longer latencies compared to B6129 controls (p = 0.0036). At the pre-symptomatic stage (3 months of age), 3xTg-AD controls did not show significant differences in escape latencies compared to B6129 controls (p = 0.2153). Two-way ANOVA with Tukey’s post hoc multiple comparisons of the main column means. All data represent mean ± SEM **p < 0.01. Sample size is indicated in parentheses next to each experimental group label.

In Figure 5, Western blot analysis of the behaviorally tested mice revealed persistent alterations in APP CTFs levels in cortical and hippocampal lysates obtained from the behaviorally tested 3-month-old mice PAE-B6129 mice compared with those from age-matched B6129 control cohorts. Specifically, PAE-B6129 mice exhibited significantly elevated levels of APP CTFs in both the cortex, up to 41.26% (p = 0.0065, unpaired Student’s t-test, Figs. 5A and 5C), and hippocampus, up to 22.62% (p = 0.0170, unpaired Student’s t-test, Figs. 5E and 5G). In contrast, the levels of PS1 CTF in PAE-B6129 showed a greater variability in the cortex (Coefficient of variation (CoV): B6129 = 5.147% vs. PAE-B6129 = 15.61%; p = 0.0725, unpaired Student’s t-test, Figs. 5A and 5C) and hippocampus (p = 0.4578, unpaired Student’s t-test, Figs. 5E and 5G) that did not differ significantly from those of the B6129 control cohort.

**Figure 5.**
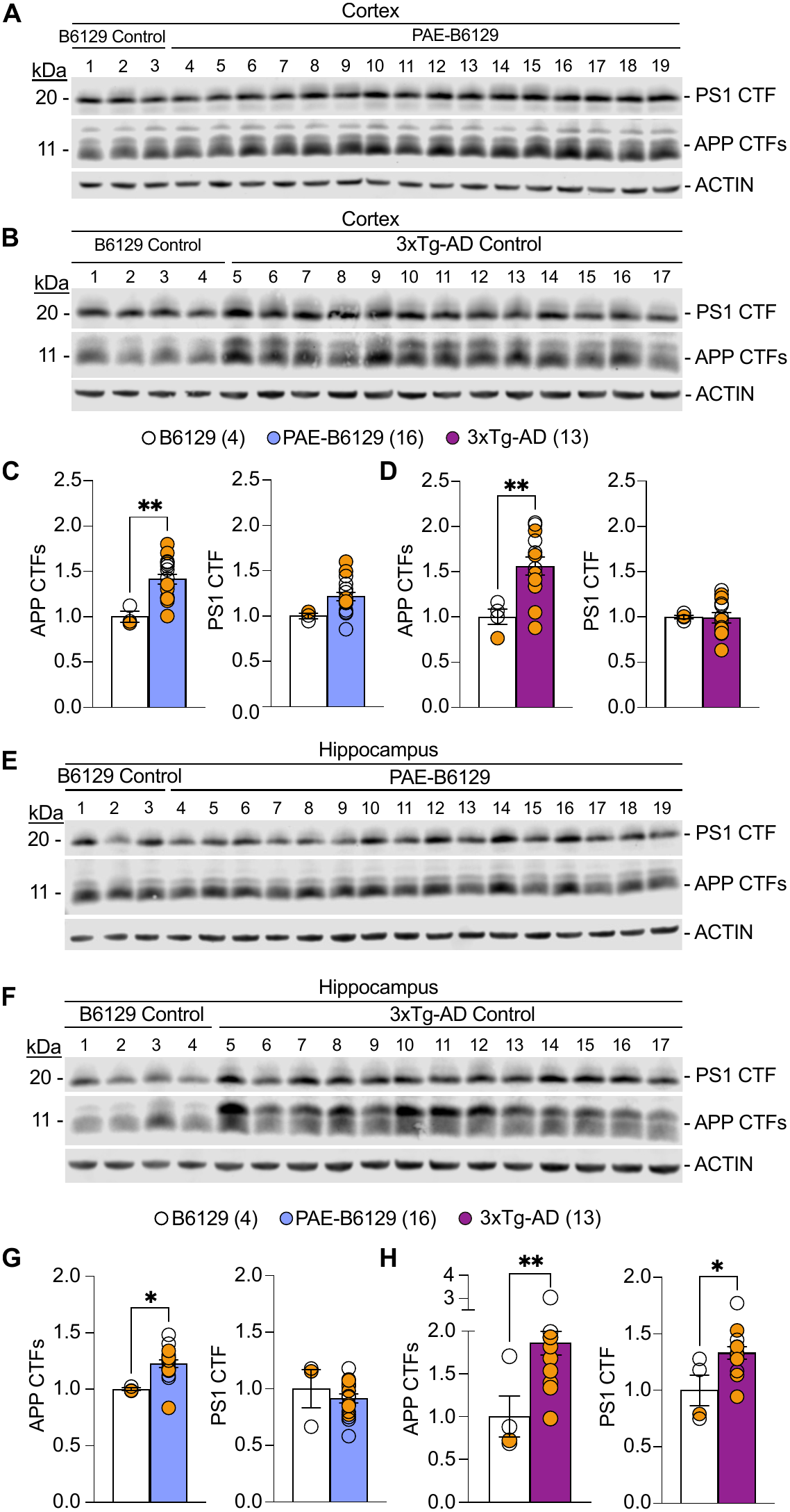
PAE reduces γ-secretase activity in B6129 mice at 3 months of age. **A)** Western analysis of cortical lysates from B6129 controls (lanes 1-3) and PAE-B6129 (lanes 4-19) mice at 3 months of age shows no change in PS1 CTF levels between groups and increased levels of APP CTFs in PAE-B6129 mice compared to B6129 controls. **(B)** Western blot analysis of cortical lysates from B6129 controls (lanes 1-4) and 3xTg-AD controls (lanes 5-17) mice at 3 months of age shows no change in levels of PS1 CTF and increased levels of APP CTFs in 3xTg-AD mice compared to B6129 controls. **(C-D)** Quantification of cortical levels of APP CTFs and PS1 CTF shows that **(C)** levels of APP CTFs are significantly increased in the cortex of PAE-B6129 (p = 0.065) compared to B6129 controls, and levels of PS1 CTF show no significant differences between PAE-B6129 (p = 0.0725) and B6129 controls. **(D)** Levels of APP CTFs (p = 0.0097) and PS1 CTF (p = 0.0165) in 3xTg-AD cortex are significantly increased compared to B6129 controls. Unpaired Student’s t-test. **(E)** Western analysis of hippocampal lysates from B6129 controls (lanes 1-3) and PAE-B6129 (lanes 4-19) show no changes in PS1 CTF level, and increased levels of APP CTFs in PAE-B6129 compared to B6129 controls. **(F)** Western analysis of hippocampal lysates from B6129 controls (lanes 1-4) and 3xTg-AD controls (lanes 5-17) shows increased levels of PS1 CTF and APP CTFs in 3xTg-AD mice compared to B6129 controls. **(G-H)** Quantification of hippocampal levels of APP CTFs and PS1 CTF shows that **(G)** levels of APP CTFs in PAE-B6129 (p = 0.0170) are significantly increased compared to B6129 controls. Levels of PS1 CTF were not significantly different between groups (p = 0.4578). **(H)** Levels of APP CTF in 3xTg-AD controls (p = 0.0096) were significantly higher compared to B6129 controls. Levels of PS1 CTF in 3xTg-AD (p = 0.0165) were significantly higher compared to B6129 controls. Unpaired Student’s t-test. All data represent mean ± SEM. *p < 0.05, **p < 0.01. Sample size is indicated in parentheses next to each experimental group label. Orange-filled and open circles represent data obtained from individual male and female mice, respectively.

Cortical and hippocampal lysates from 3-month-old 3xTg-AD mice without PAE, which served as pathological controls in the Morris Water Maze test, were also analyzed by Western blotting (Fig. 5). In the cortex, PS1 CTF levels from 3xTg-AD mice did not differ from those in B6129 controls (p = 0.9235, unpaired Student’s t-test, Figs. 5B and 5D). However, in the hippocampus, PS1 CTF levels were significantly increased up to 33.12% (p = 0.0165, unpaired Student’s t-test; Figs. 5F and 5H), likely reflecting overexpression of the human PS1 M146V transgene. By contrast, APP CTFs were significantly increased in 3xTg-AD mice in both the cortex, up to 56.22% (p *=* 0.0097, unpaired Student’s t-test; Figs. 5B and 5D) and hippocampus, up to 86.41% (p *=* 0.0096, unpaired Student’s t-test; Figs. 5F and 5H). When these increases were compared with APP CTF levels observed in PAE-B6129 mice, the data suggest that both prenatal alcohol exposure and the PS1 M146V mutation produce a comparable reduction in γ-secretase activity in young 3-month-old adult mice.

To determine whether the behavioral and γ-secretase alterations observed at 3 months of age persist or worsen with age, we next examined hippocampal-dependent learning using the same regimen illustrated in Figure 4, and γ-secretase–related processing in PAE B6129 mice at 6 months of age (Figs. 6 and 7). Figure 6 illustrates that 6-month-old PAE-B6129 mice exhibited more severe deficits in hippocampal-dependent learning. During hidden platform training, PAE-exposed mice displayed highly significant longer escape latencies compared with age-matched non-PAE B6129 controls (p < 0.0001, two-way ANOVA with Tukey’s post hoc multiple comparisons of the main columns means), and the differences in escape latencies between PAE-B6129 and 3xTg-AD controls mice are relatively modest (p = 0.0220, two-way ANOVA with Tukey’s post hoc multiple comparisons of the main columns means).

**Figure 6.**
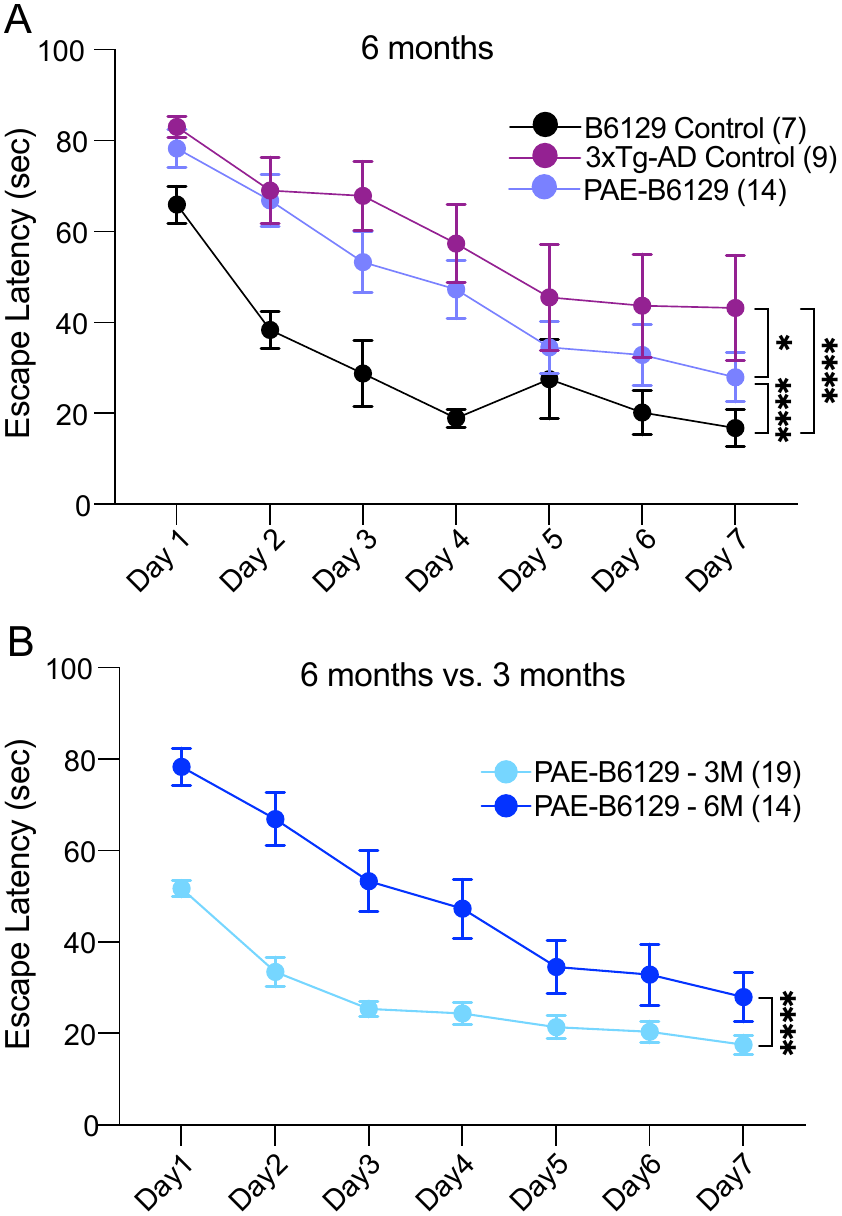
PAE impairs learning and memory and γ-secretase activity in B6129 mice at 6 months of age. **(A)** During the hidden platform version of the Morris water maze test, PAE-B6129 mice at 6 months of age display significantly longer escape latencies compared to B6129 controls (p < 0.0001). Similarly, 3xTg-AD controls (p < 0.0001) show significant differences in escape latencies compared to B6129 controls. The difference in escape latency between PAE-B6129 and 3xTg-AD mice reached modest statistical significance (p = 0.0220). (B) Comparison of 3- and 6-month-old PAE-B6129 mice revealed significantly longer latencies in older mice (p < 0.0001), consistent with an age-associated progression of learning and memory deficits following PAE. Two-way ANOVA with Tukey’s post hoc multiple comparisons of the main columns means. All data represent mean ± SEM *p < 0.05 ****p < 0.0001. Sample size is indicated in parentheses next to each experimental group label.

**Figure 7.**
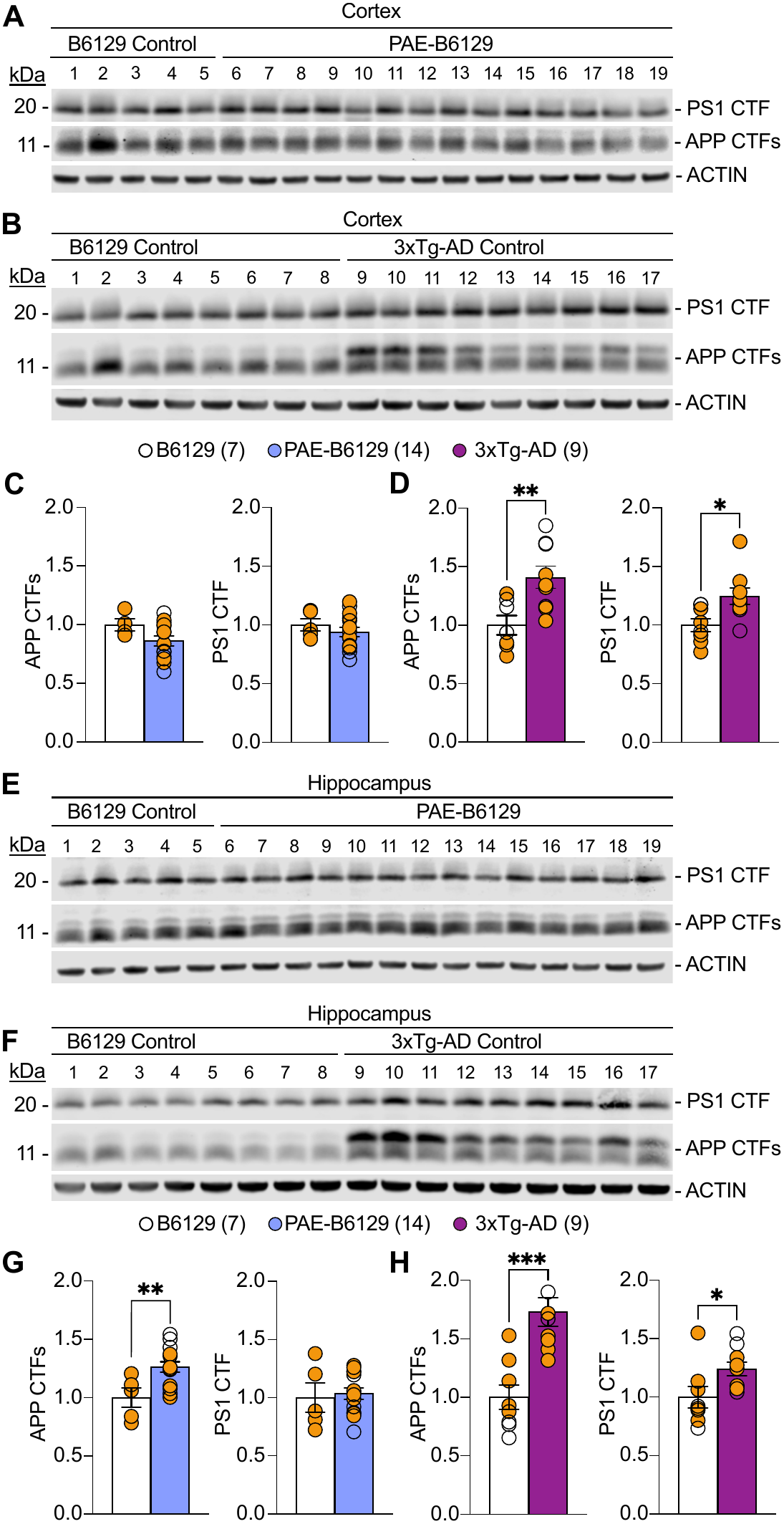
PAE reduces γ-secretase activity in the hippocampus of B6129 mice at 6 months of age. **(A)** Western analysis of cortical lysates from B6129 controls (lanes 1-5) and PAE-B6129 (lanes 6-19) mice at 6 months of age shows no change in PS1 CTF and APP CTFs levels between groups **(B)** Western blot analysis of cortical lysates from B6129 controls (lanes 1-4) and 3xTg-AD controls (lanes 5-17) mice at 6 months of age shows no change in levels of PS1 CTF and increased levels of APP CTFs in 3xTg-AD mice compared to B6129. **(C-D)** Quantification of cortical levels of APP CTFs and PS1 CTF shows that **(C)** levels of APP CTFs (p = 0.1219) and PS1 CTF (p = 0.4461) are not significantly different in the cortex of PAE-B6129 compared to non-alcohol-exposed B6129 **(D)** Levels of APP CTFs (p = 0.0067) and PS1 CTF (p = 0.0165) in 3xTg-AD cortex are significantly increased compared to B6129. Unpaired Student’s t-test. **(E)** Western analysis of hippocampal lysates from B6129 controls (lanes 1-3) and PAE-B6129 (lanes 4-19) shows no changes in PS1 CTF level, and increased levels of APP CTFs in PAE-B6129 compared to B6129 controls. **(F)** Western analysis of hippocampal lysates from B6129 controls (lanes 1-4) and 3xTg-AD controls (lanes 5-17) shows increased levels of PS1 CTF and APP CTFs in 3xTg-AD mice compared to B6129 controls. **(G-H)** Quantification of hippocampal levels of APP CTFs and PS1 CTF shows that **(G)** levels of APP CTFs in PAE-B6129 (p = 0.0094) are significantly increased compared to B6129 controls. Levels of PS1 CTF were not significantly different between groups (p = 0.7407). **(H)** Levels of APP CTF in 3xTg-AD controls (p = 0.0004) are significantly higher compared to B6129 controls. Levels of PS1 CTF in 3xTg-AD (p = 0.0393) are significantly higher compared to B6129 controls. Unpaired Student’s t-test. All data represent mean ± SEM. *p < 0.05, **p < 0.01, ***p < 0.001. Sample size is indicated in parentheses next to each experimental group label. Orange-filled and open circles represent data obtained from individual male and female mice, respectively.

To further investigate the age-related progression of learning and memory deficits following PAE, we directly compared escape latencies in 3- and 6-month-old B6129 control mice. Older mice exhibited significantly longer escape latencies than younger mice (p < 0.0001, two-way ANOVA with Tukey’s post hoc multiple comparisons of the main columns means), indicating impaired performance in the Morris water maze with increasing age. This indicates that spatial learning impairments not only persisted but also worsened into mid-adulthood, reaching a level comparable to that observed in the 3xTg-AD mouse model.

In cortical and hippocampal lysates from 6-month-old behaviorally tested mice, PAE-B6129 mice showed no significant changes in APP CTFs levels in the cortex compared with non-exposed B6129 controls (p = 0.1219, unpaired Student’s t-test; Figs. 7A and 7C). In contrast, APP CTF levels in the hippocampus displayed a significant increase in PAE-B6129 mice up to 26.16% (p = 0.094, unpaired Student’s t-test; Fig. 7E and 7G), which can be related to the learning and memory deficits observed in this group. Consistent with earlier observations, PS1 CTF levels did not differ significantly in cortex (p = 0.4461, unpaired Student’s t-test) and hippocampus (p = 0.7407, unpaired Student’s t-test) compared to the B6129 control group, indicating that the catalytic subunit of the γ-secretase complex remains unchanged (Figs. 7C and 7G). Analysis of APP CTF levels in the 3xTg-AD mouse model revealed a significant increase in the cortex up to 40.93% (p = 0.0067, unpaired Student’s t-test; Figs. 7B and 7D), with an even greater increase in the hippocampus, up to 73.01% (p = 0.0004, unpaired Student’s t-test; Figs. 6F and 6H), supporting a potential link between more pronounced γ-secretase deficits in hippocampal regions and learning and memory impairments. As observed in 3-month-old mice (Fig. 5), 3xTg-AD mice exhibited higher PS1 CTF protein levels in the cortex (24.39%, p = 0.0162, unpaired Student’s t-test; Fig. 7D) and hippocampus (24.07%, p = 0.0393, unpaired Student’s t-test; Fig. 7H) compared to B6129 controls, consistent with overexpression of the PS1 M145V transgene. Together, these findings suggest that early PAE-induced disruption of γ-secretase–mediated APP processing produces long-lasting brain region-selective alterations. In particular, the pronounced accumulation of APP CTFs observed in the cerebral cortex at birth and in the hippocampus at adulthood of PAE-B6129 mice strongly suggests a mechanistic link to the persistent hippocampal-dependent learning deficits observed in 3 and 6-month-old mice.

### PAE alters learning and memory in 3xTg-AD mice

To investigate whether PAE contributes to pathological outcomes in mice carrying AD–linked mutations, we asked whether PAE exacerbates learning and memory deficits in 3xTg-AD mice at an early stage of the disease (4-month-old) and, if so, whether it is associated with alterations in γ-secretase–dependent APP processing. Hippocampal-dependent learning and memory were assessed in PAE and non-exposed 3xTg-AD mice (Fig. 8A). As expected, non-exposed 3xTg-AD mice already displayed early impairments relative to B6129 controls by 4 months of age (p = 0.0299, two-way ANOVA with Tukey’s post hoc multiple comparisons of the main columns means), consistent with baseline deficits driven by PS1, APP, and Tau mutations. PAE-3xTg-AD mice exhibited further delays in escape latency in the Morris water maze compared to non-exposed 3xTg-AD mice, with escape latency significantly greater than B6129 controls (p = 0.0005, two-way ANOVA with Tukey’s post hoc multiple comparisons of the main columns means), indicating that early-life alcohol exposure amplifies pre-existing spatial learning deficits. Although learning and memory impairments are only beginning to emerge at 4 months of age, PAE-3xTg-AD mice already exhibit more pronounced deficits than their non-exposed 3xTg-AD counterparts, suggesting that PAE may accelerate the emergence and increase the severity of AD-associated behavioral dysfunction. We anticipate that these differences in learning and memory performance may become more pronounced as the mice age and AD-related pathology progress.

**Figure 8.**
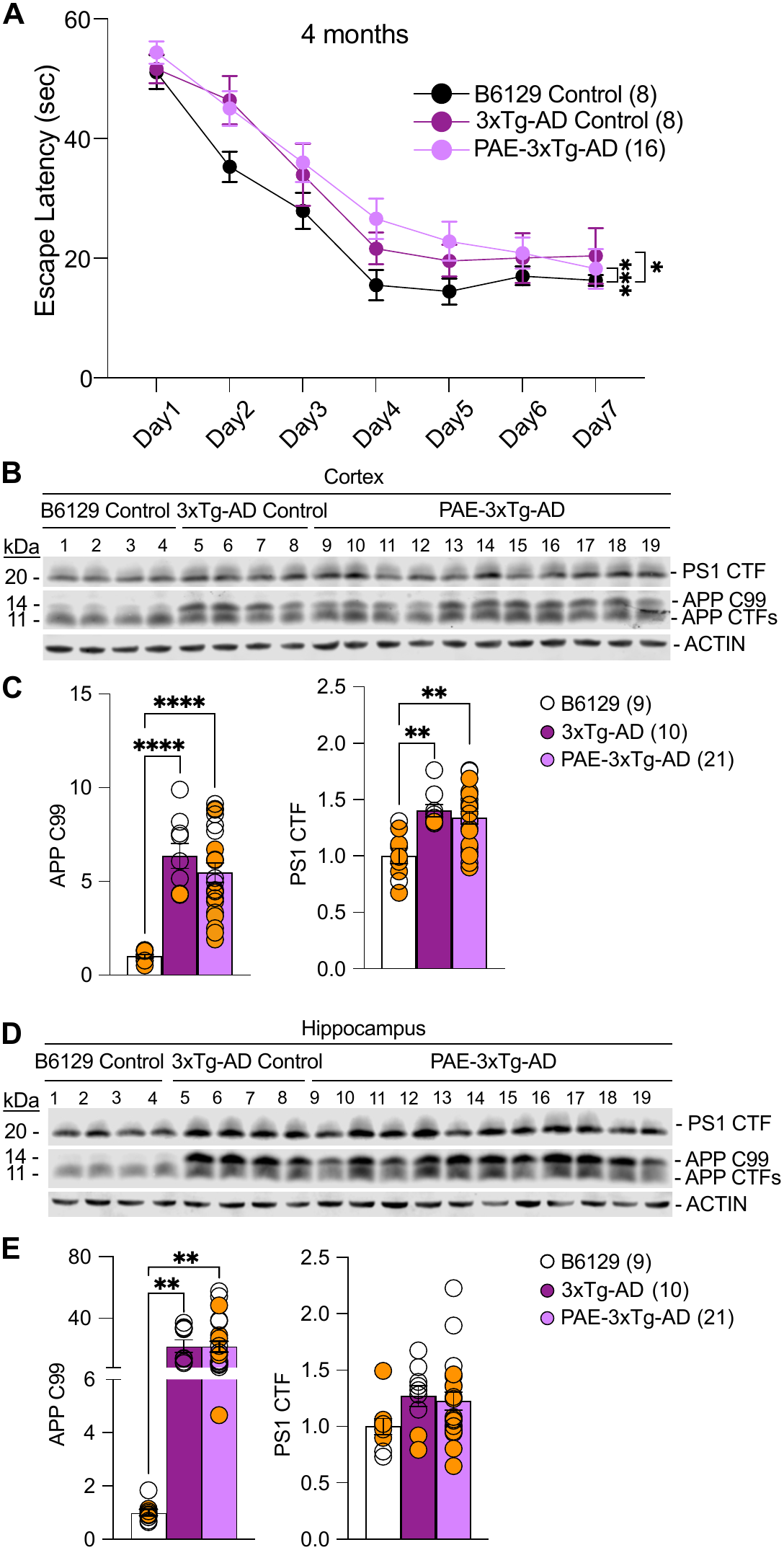
PAE alters learning and memory in 3xTg-AD mice in the pre-symptomatic stage. **(A)** During the hidden platform version of the Morris water maze test, PAE-3xTg AD mice at 4 months of age (pre-symptomatic stage) display significantly longer escape latencies compared to B6129 controls (p = 0.0005). 3xTg-AD controls (p = 0.0299) show less significant differences in escape latencies compared to B6129 controls. Two-way ANOVA with Tukey’s post hoc multiple comparisons of the main columns means. All data represent mean ± SEM *p < 0.05 ***p < 0.001. Sample size is indicated in parentheses next to each experimental group label. Orange-filled and open circles represent data obtained from individual male and female mice, respectively.

Western blot analyses of γ-secretase activity in cortical (Fig. 8B, 8C) and hippocampal (Fig. 8D, 8E) lysates, from the same behaviorally tested mice, showed increased levels of APP CTFs, with a more pronounced increase in APP C99, in both PAE and non-exposed 3xTg-AD mice compared with B6129 controls, consistent with the effects observed in PS1 mutant models. Significant overexpression of the PS1 M146V transgene was detected in cortical tissue from non-exposed 3xTg-AD (40.18%, p *=* 0.0019, one-way ANOVA with Tukey’s post hoc multiple comparisons) and PAE-3xTg-AD mice (33.87%, p *=* 0.0020, one-way ANOVA with Tukey’s post hoc multiple comparisons) relative to the B6129 control group (Fig. 8C). In the hippocampus, PS1 CTF levels were elevated but did not reach statistical significance (3xTg-AD, 26.82%, p *=* 0.1928, and PAE-3xTg-AD, 22.53%, p *=* 0.1966, one-way ANOVA with Tukey’s post hoc multiple comparisons), likely reflecting greater inter-animal variability in transgene expression and/or differential regional susceptibility to PAE (Fig. 8E).

In both cortex and hippocampus, protein analysis revealed a significant increase in APP C99 levels in the cortex, 6 times higher (Figs. 8B and 8C, 3xTg-AD controls, p *<* 0.0001; PAE-3xTg-AD, *p* < 0.0001, one-way ANOVA with Tukey’s post hoc multiple comparisons) and hippocampus, 20 times higher (Figs. 8D and 8E, 3xTg-AD controls, p *=* 0.0063; PAE-3xTg-AD, p *=* 0.0012, one-way ANOVA with Tukey’s post hoc multiple comparisons) compared with the B6129 control group. APP C99 levels did not differ significantly between 3xTg-AD and PAE-3xTg-AD mice; however, greater variability was observed within the PAE group (CoV: cortex 3xTg-AD vs. PAE-3xTg-AD = 31.70% vs. 42.71%; hippocampus 3xTg-AD vs. PAE-3xTg-AD = 55.47% vs. 75.01%), which may reflect differences in individual vulnerability to prenatal alcohol exposure, variability in transgene expression, or an interaction between both factors.

## Discussion

Preclinical evidence suggests that PAE may be a potential early-life risk factor that predisposes the brain to neurodegenerative vulnerability later in life, including susceptibility to AD/ADRD (Alhowail, 2022; Gerlikhman et al., 2023; Tousley et al., 2023; Walter et al., 2023). However, there are currently no human epidemiological datasets to substantiate this preclinical concept. One reason may be that FASD was not formally identified as a neurodevelopmental condition until the late 1960s (Lemoine, 1968; Jones et al., 1973; Jones & Smith, 1973). As a result, the earliest individuals who were identified under the FASD framework are only now approaching the age at which AD/ADRD pathology and clinical symptoms typically begin to emerge. Consequently, there are no records of clinically diagnosed FASD individuals who have been systematically evaluated for Alzheimer’s disease pathology or dementia, creating an opportunity for clinical and epidemiological studies to determine whether early-life alcohol exposure confers long-term risk for age-related neurodegeneration. In this context, preclinical studies are essential for establishing the biological basis for PAE as a developmental contributor to AD/ADRD risk.

The present study provides evidence that PAE disrupts γ-secretase–related processing during brain development, and that these alterations are associated with persistent deficits in hippocampal-dependent learning and memory later in life, reminiscent of AD/ADRD. Using a binge-like gestational exposure paradigm, we demonstrate that PAE induces early accumulation of APP CTFs and alters Notch-related processing in the short term that persist into adulthood and are accompanied by behavioral impairments, consistent with the hypothesis that PAE establishes long-lasting neurobiological changes that increase vulnerability to learning and memory decline.

It is noteworthy that the selection of ages for this study captures both pre-symptomatic and early symptomatic stages of pathology. In the 3xTg-AD mouse model, subtle synaptic dysfunction emerges as early as 3–4 months of age, preceding cognitive decline (Oddo et al., 2003). This pre-symptomatic window provides an opportunity to interrogate early alterations, particularly those related to γ-secretase activity and memory loss, before the onset of confounding widespread neurodegeneration. Incorporating age-matched wild-type cohorts allows for the detection of subtle, early memory impairments. The presence of early deficits in wild-type mice is especially relevant in the context of PAE, where neurodevelopmental insults may shift the baseline trajectory of cognitive function independent of AD transgenes. Assessing animals at 3, 4, and 6 months of age captures the progression from pre-symptomatic to more advanced stages, enabling differentiation between early vulnerability and later-stage pathology.

A key finding of this study is that a brief maternal binge-like ethanol exposure during mice gestational days 13–15, a window that approximates the late first trimester to early second trimester of human gestation (Peavey & Dotters-Katz, 2021) , and is characterized by robust neurogenesis, neuronal migration, and early cortical and hippocampal circuit formation (Hill, 2023; Otis & Brent, 1954; Parnell et al., 2014), is sufficient to disrupt γ-secretase–dependent APP processing in the neonatal brain. The accumulation of APP CTFs in both cortex and hippocampus suggests reduced γ-secretase activity, as impaired cleavage of APP C99 is a recognized biochemical indicator of γ-secretase dysfunction (Bourgeois et al., 2018; Pulina et al., 2020). These effects were observed not only in 3xTg-AD mice carrying AD-linked mutations but also in wild-type mice, indicating that alcohol exposure alone can induce a molecular phenotype resembling presenilin-related dysfunction. This finding provides a potential mechanistic framework linking developmental alcohol exposure to pathways implicated in neurodegenerative disease.

The increased accumulation of APP CTF in cortical layers II–VI and hippocampal CA subregions is particularly relevant given the known vulnerability of these regions to alcohol-induced developmental insult (Cuzon et al., 2008; Delatour et al., 2019, 2020; Skorput & Yeh, 2016). Previous work has shown that PAE disrupts neuronal migration, dendritic maturation, and synaptic development in the cortex and hippocampus (Berman & Hannigan, 2000; Delatour et al., 2019, 2020; Guerri, 2002; Livy et al., 2003; Miller, 1993; Subbanna & Basavarajappa, 2022). Our findings suggest that impaired γ-secretase signaling may contribute to these abnormalities by altering developmental pathways regulated by APP and Notch processing.

The PAE effect on γ-secretase–related processing of APP persisted into adulthood. At both 3 and 6 months of age, PAE wild-type mice exhibited impaired performance in the Morris water maze, accompanied by persistent accumulation of APP CTFs in the hippocampus. These data suggest that developmental alcohol exposure produces long-lasting alterations in protein processing pathways that may contribute to sustained hippocampal dysfunction. The progressive worsening of escape latency by 6 months further supports the notion that early molecular injury induced by alcohol exposure has enduring functional consequences. The relationship between these phenotypes is not necessarily linear across age: although PAE produced a more pronounced learning deficit at 6 months in B6129 mice, the magnitude of APP CTF accumulation was greater at the earlier time point. Importantly, we consider the possibility that the greater APP CTF accumulation observed at earlier ages may represent an early molecular insult whose functional consequences become evident later in life. In this context, the temporal dissociation between the molecular and behavioral phenotypes could be consistent with a “two-hit” model, in which an early-life insult induced by PAE creates or primes a pathological vulnerability that subsequently manifests as cognitive dysfunction with aging (Gunn et al., 2019).

Morris Water Maze performance reflects multiple cognitive and behavioral processes and should not be interpreted solely as a generalized measure of “memory impairment.” In the present study, hidden platform acquisition trials primarily assessed hippocampal-dependent spatial learning. PAE mice did not exhibit deficits during visible platform testing, suggesting that the observed impairments were unlikely to result from visual or motor dysfunction. These findings support the interpretation that PAE correlates with a disruption of hippocampal-dependent spatial learning and memory processes in adulthood.

The findings in 3xTg-AD mice provide additional evidence that PAE may influence outcomes in the context of pre-existing genetic vulnerability. PAE was associated with greater learning deficits in 4-month-old transgenic mice, an age at which early cognitive abnormalities begin to emerge in this model (Bourgeois et al., 2018; Javonillo et al., 2022). Although APP C99 accumulation was already elevated due to AD-associated mutations, the greater behavioral impairment observed following PAE suggests that prenatal alcohol exposure may further contribute to cognitive deficits in the presences of AD-associated genetic risk.

The absence of PAE-induced changes in PS1 CTF levels suggests that prenatal alcohol exposure does not directly affect the abundance of the presenilin protein. Rather, these findings support the presence of a regulatory mechanism acting at the level of γ-secretase function. Such regulation may involve altered catalytic efficiency, impaired complex maturation, changes in subcellular localization, or modifications in the interaction between γ-secretase and its substrates. Together, this suggests that PAE may influence APP processing through functional impairment of the proteolytic machinery rather than through altered expression of its core components. In summary, the plots presented in Figures 5, 7 and 8, males (orange circles) and females (clear circles) were similarly distributed across experimental groups in both the APP C-terminal fragment and PS1 C-terminal fragment analyses. This suggests that sex did not substantially influence the effects of PAE on γ -secretase activity, and that PAE-associated alterations in APP processing were generally consistent across sexes. However, because these studies were not designed or powered to detect sex-specific effects, further work is needed to determine whether PAE produces sex-dependent outcomes. The findings of the present study bear important implications, given that binge drinking during early pregnancy is clinically relevant, particularly during periods when pregnancy may not yet have been recognized - even a brief exposure during a relatively narrow developmental window may produce persistent molecular alterations in pathways critical for brain maturation and cognitive function later in life. This supports the concept that PAE may not only contribute to neurodevelopmental disorders such as FASD but may also increase vulnerability to age-related neurodegenerative processes.

Several limitations should be acknowledged. While the accumulation of APP CTFs strongly supports impaired γ-secretase activity, this interpretation is based on indirect biochemical readouts. Direct quantification of γ-secretase cleavage products, including Aβ40, Aβ42, and AICD, will be important in future studies to more precisely define how PAE alters enzyme activity, substrate processing, and peptide production. Assessing Aβ42/Aβ40 ratios will determine whether PAE shifts γ-secretase cleavage toward more aggregation-prone species, a key feature of AD/ADRD pathology.

In addition, the present study focuses on early and mid-adulthood time points and therefore does not address whether PAE accelerates the progression of AD-like neuropathology in older mice. Extended longitudinal studies will be necessary to associate early-life disruption of γ-secretase function with enhanced amyloid deposition, tau pathology, or progressive neuronal loss later in life. Future work should incorporate comprehensive neuropathological analyses, including assessment of amyloid plaque burden, tau hyperphosphorylation, synaptic function, and region-specific neurodegeneration in vulnerable brain areas such as the hippocampus and cortex.

While our behavioral and biochemical findings suggest persistent hippocampal dysfunction, direct evaluation of neurodegenerative processes, such as neuronal loss, dendritic spine alterations, and neuroinflammatory responses, was not performed. Given that accumulation of APP CTFs, particularly C99, has been implicated in synaptic toxicity and endosomal dysfunction (Bourgeois et al., 2018), it will be important to determine whether PAE-induced alterations in APP processing contribute to progressive synaptic degeneration and circuit-level dysfunction. Assessing markers of gliosis and neuroinflammation will also be essential to establish whether PAE primes the brain for heightened inflammatory responses that may exacerbate neurodegenerative processes over time.

The variability observed in PAE cohorts may reflect differences in transgene expression, differential susceptibility to PAE, or interactions between genetic background and environmental exposure (Chen et al., 2011; Everson & Eberhart, 2022). These findings open avenues for future research aimed at elucidating the molecular mechanisms underlying individual variability in vulnerability to PAE-induced effects.

Future investigations should systematically assess how the timing, duration, and developmental stage of prenatal alcohol exposure (PAE) influence γ-secretase regulation, APP processing, and, commensurately, learning and memory. Direct comparisons with acute exposure paradigms, such as the 3-day binge drinking model, will be needed to determine whether the observed molecular alterations reflect cumulative developmental reprogramming or transient ethanol-induced perturbations. In addition, PAE at different gestational time points should be evaluated, as the embryonic and fetal brain undergoes stage-specific waves of neurogenesis, synaptic formation, and circuit refinement that may differentially determine vulnerability to long-term Alzheimer’s disease–related molecular and neurobehavioral changes.

Although PAE did not result in a statistically significant exacerbation of the behavioral deficits observed in 3xTg-AD mice compared with their non-exposed counterparts, this finding should be interpreted with caution. The relatively small sample size may have limited the statistical power to detect more subtle differences between these groups. Notably, the PAE-3xTg-AD mice exhibited a trend toward greater behavioral impairment, suggesting a potential worsening of the phenotype following prenatal exposure. Thus, while the current data do not provide sufficient statistical evidence to conclude that PAE exacerbates the learning and memory deficits associated with the 3xTg-AD genotype, the observed trend warrants further investigation in larger cohorts and at later ages, when AD-related behavioral deficits are expected to become more pronounced.

In summary, our findings identify disrupted γ-secretase–related processing as a novel molecular consequence of PAE and support a mechanistic link between early-life binge alcohol exposure and persistent hippocampal-dependent learning and memory dysfunction. These data provide the first experimental evidence that developmental alcohol exposure impacts core proteolytic pathways central to AD/ADRD pathogenesis.

## Materials and Methods

### Wild-type and Transgenic Mice

Genotype-verified breeder pairs were obtained directly from The Jackson Laboratory. Wild-type B6129SF1/J (B6129) Strain # 101043 (https://www.jax.org/strain/101043) and B6;129-Tg (APPSwe, tauP301L)1Lfa *Psen1^tm1Mpm^*/Mmjax (3xTg-AD) MMRRC Strain # 034830-JAX (https://www.jax.org/strain/004807) were bred and maintained following guidelines and procedures approved by the Institutional Animal Care and Use Committee (IACUC) at Dartmouth College. Animals were housed in a 12 h light/dark cycle with light periods occurring between 7:00 AM and 7:00 PM.

No animal attrition occurred within the experimental groups during the study, and the number of animals assigned to each group at the beginning of the experimental period remained unchanged through the completion of the study. However, reduced litter size was observed following prenatal alcohol exposure (PAE). In the 3xTg-AD colony, control litters typically consisted of approximately eight pups, whereas PAE litters occasionally contained only four to six pups. Thus, the lower number of offspring available for the study reflects reduced litter size following PAE rather than animal loss or attrition during the experimental period.

### Prenatal Alcohol Exposure (PAE)

As previously described, in our binge-type alcohol exposure paradigm illustrated in Figure 1A (Delatour et al., 2019, 2020; Skorput & Yeh, 2016; Tousley et al., 2023), breeding trios of one male and two female mice were housed overnight for breeding, with the following day designated as embryonic day 0.5 (E0.5) and after which dams were singly housed and acclimated to an *ad libitum* control liquid diet from E10 to E12. From E13 to E15, feeding bottles were replaced daily with either 5% (w/w) ethanol liquid diet or an isocaloric control liquid diet (Table 1), with water available *ad libitum*. Dams were weighed daily throughout the PAE period. Wild-type B6129 dams reached a mean blood ethanol concentration (BEC) of 0.20 g/dL ± 0.006 g/dL, whereas 3xTg-AD dams reached a mean BEC of 0.11 g/dL ± 0.005 g/dL, as measured at 11:00 PM on E15.5 using an Analox AM1 series III analyzer (Analox Instruments). The lower BEC observed in 3xTg-AD dams compared with B6129 dams may reflect differences in ethanol preference, sensitivity, or metabolic processing between the two genotypes, as daily liquid diet intake exceeded 10 g in both B6129 (Figure 1B) and 3xTg-AD (Figure 1C) throughout the acclimation and experimental periods (Lim et al., 2012; Rhodes et al., 2005).Importantly, both groups achieved BECs exceeding 0.08 g/dL, thereby meeting the NIAAA criteria for binge-level alcohol exposure (NIAAA, 2025).

**Table 1.** Isocaloric liquid diet formulas for 1 liter preparation (w/w)

| **Diet ID** | **EtOH (mL)** | **Premix<sup>1</sup> (g)** | **Olive Oil (g)** | **Maltose<sup>2</sup> (g)** | **ddH2O (mL)** |
|---|---|---|---|---|---|
| Control | 0 | 109 | 28.4 | 87.5 | 775.1 |
| 5% EtOH | 65 | 109 | 28.4 | 0 | 812.6 |

1. Premix L10251A (Research Diets)
2. Maltose Dextrin 42 I20005 (Research Diets)

The dams used in this study were young adult females (2 to 3 months of age) at the time of breeding. Because chronological age does not translate linearly between mice and humans, particularly during development and reproductive maturation, we have avoided assigning a precise human-age equivalent. Based on established comparative developmental frameworks and calculations, these animals represent approximately 20-year-old young adult females at the height of their reproductive stage, rather than a more advanced child-bearing age (Dutta & Sengupta, 2016).

Age-matched mice fed with the control liquid diet were used as non-ethanol-exposed control groups for all experimental comparisons. Non-ethanol-exposed B6129 mice served as genotype-matched controls for the PAE-B6129 cohort. Similarly, non-exposed 3xTg-AD mice served as baseline transgenic controls, providing the reference for AD-like pathological profile. This cohort served not only as the direct control for the PAE-3xTg-AD group, but also as an AD-relevant benchmark for comparison with the PAE-B6129 group, facilitating evaluation of whether PAE alone induces molecular changes that approach those observed in the 3xTg-AD model.

### Western Blot Analysis

After birth, brains from P0 mice were harvested, whereas the remaining experimental mice were maintained for subsequent behavioral and biochemical analyses at 3, 4, and 6 months of age. The entire right hemisphere was reserved for immunohistochemical analysis. From the left hemisphere, the whole cortex was dissected from P0 mice, while the cerebral cortex and hippocampus were separately dissected at 3, 4, and 6 months of age. All tissues were fast frozen and stored for subsequent protein lysate preparation. Cortical and hippocampal tissues were manually homogenized using pestle homogenizers followed by passage through 1ml syringes fitted with 25G needles in radioimmunoprecipitation assay buffer, Pierce RIPA buffer (ThermoFisher Scientific, Cat No. PI89900) supplemented with phenylmethylsulfonyl fluoride (PMSF) protease inhibitor (Sigma, Cat No. 10837091001) and Halt protease inhibitor cocktail, EDTA-free (ThermoFisher Scientific, Cat No. 78425). Protein concentrations were quantified using the Pierce BCA protein assay kit (ThermoFisher Scientific, Cat No. 23225). Protein lysates were normalized to 3 ug/ul in NuPAGE LDS sample buffer (ThermoFisher, Cat No. NP0007) and NuPAGE sample reducing agent (ThermoFisher, Cat No. NP0009). Protein samples (20 - 40 μg total protein) were resolved on NuPAGE 4-12% Bis-Tris midi SDS-PAGE gels (ThermoFisher Scientific, Cat No. WG1402BOX) and transferred to a 0.45 μm nitrocellulose membrane (ThermoFisher Scientific, Cat. No. 88018) using wet transfer conditions. Membranes were blocked for 1 h in Intercept (TBS) blocking buffer (LICORbio, Cat. No. 927-60001) and incubated overnight at 4°C with primary antibodies specific for PS1 CTF, APP CTFs, and NICD (Table 2). Detection was performed using infrared-conjugated secondary antibodies (Table 2) and the Odyssey CLx imaging system (LICORbio). Protein bands were quantified with Fiji image processing software (ImageJ), with target protein levels normalized to actin as a loading control.

**Table 2.**
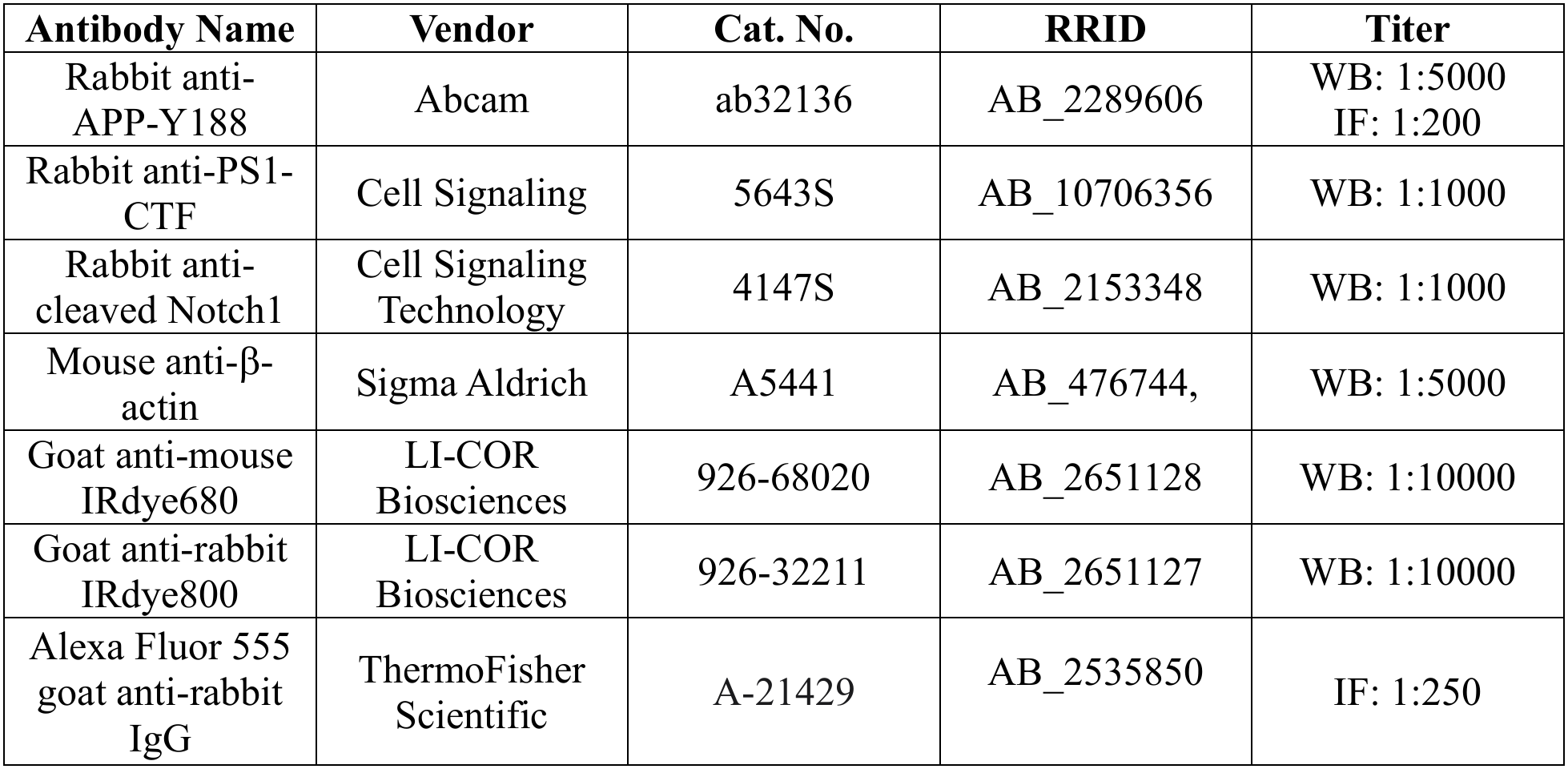
List of antibodies used in this study.

| **Antibody Name** | **Vendor** | **Cat. No.** | **RRID** | **Titer** |
|---|---|---|---|---|
| Rabbit anti-APP-Y188 | Abcam | ab32136 | AB_2289606 | WB: 1:5000<br>IF: 1:200 |
| Rabbit anti-PS1-CTF | Cell Signaling | 5643S | AB_10706356 | WB: 1:1000 |
| Rabbit anti-cleaved Notch1 | Cell Signaling Technology | 4147S | AB_2153348 | WB: 1:1000 |
| Mouse anti-β-actin | Sigma Aldrich | A5441 | AB_476744, | WB: 1:5000 |
| Goat anti-mouse IRdye680 | LI-COR Biosciences | 926-68020 | AB_2651128 | WB: 1:10000 |
| Goat anti-rabbit IRdye800 | LI-COR Biosciences | 926-32211 | AB_2651127 | WB: 1:10000 |
| Alexa Fluor 555 goat anti-rabbit IgG | ThermoFisher Scientific | A-21429 | AB_2535850 | IF: 1:250 |

### Behavioral Analysis

Mice were individually handled daily for 3 days, 5 min each, before testing. Three cohorts of naïve mice: Cohort 1: ∼2.5 months of age; wild-type B6129 control (N=9), 3xTg-AD control (N=15), PAE-B6129 (N=19), Cohort 2: ∼5.5 months of age; B6129 control (N=7), 3xTg-AD control (N=9), PAE-B6129 (N=14), Cohort 3: ∼3.5 months of age; B6129 control (N=8), 3xTg-AD control (N=8), PAE-3xTg-AD (N=16). Each mouse was trained in the hidden platform version of the task for a total of 7-8 days. The Morris water maze is a circular pool of 120 cm in diameter. The release point for each training trial was varied among all four quadrants in a pseudorandom manner. Four distal visual cues were placed on the walls during training and full-cue probe tests. The escape platform (10 cm in diameter) was submerged 1 cm beneath the water surface, which was covered with plastic beads to prevent visual localization of the platform and maintained in a constant position. Mice received four training trials daily with a maximum duration of 60-90 s and an inter-trial interval of at least 15 min. Mice that did not locate the platform during a training trial were guided to the platform and allowed to remain on it for 15 s. The location of mice in the pool was continuously monitored using an automated video tracking system (Ethovision XT, Noldus). Probe trial tests were conducted on the morning of the last day (days 8-9) by removing the hidden platform and releasing mice from the opposite quadrant, allowing them to search for the platform location for 60s. To verify visual and swimming ability, the visual-cue version of the task was performed on days 8 or 9 with all distal cues removed. Four trials were administered, with the platform position marked by a visible object.

### Histological Analysis

Upon completion of behavioral testing, mice were asphyxiated with CO2 and transcardially perfused with phosphate-buffered saline solution (PBS, pH 7.4) containing 0.25 g/L heparin (Sigma, Cat. No. H3393). Brains were hemisected, and the right hemispheres were post-fixed in 4% paraformaldehyde (Sigma, Cat. No. 158127) in PBS at 4°C with gentle shaking overnight. Subsequently, the brains were embedded in paraffin following standard procedures (Kang et al., 2021). Sagittal paraffin sections (10 µm) were obtained using an Epredia HM 355 S rotary microtome. For immunofluorescence staining, paraffin sagittal sections were first deparaffinized in Histo-Clear (Mercedes Scientific, Cat. No. NAT HS202), alcohol-dehydrated in sequential baths, and then incubated in antigen retrieval buffer (10 mM Citric Acid, 0.05% Tween 20, pH 6.0) for 30 min at 95°C. After cooling down, the sections were subjected to permeabilization with Tris-buffered saline containing 0.5% Triton X-100 for 30 min on a horizontal rocker. Sections were thoroughly washed with 1X PBS solution, transferred to a humid chamber, and blocked with 10% normal goat serum (MP Biomedical, Cat. No. 219135680) for 1.5 h, at RT. Sections were incubated with primary antibody against APP CTF (Table S2) overnight at 4 °C, followed by 1.5 h RT incubation with fluorophore-conjugated Alexa 555 secondary antibody (Table S2). All sections were counterstained with DAPI (Sigma, Cat.No. D9542). Immunofluorescent images were taken and analyzed using Zeiss LSM 510 Meta laser confocal microscope (Zeiss).

### Data Quantification and Statistical Analysis

Data acquisition and quantification were performed blinded to genotype and treatment, with the exception of the Western blot analyses. All statistical analyses were performed using Prism (Version 11; GraphPad) and Excel (Microsoft). All data are presented as the mean ± SEM. The exact sample size (e.g., the number of mice) of each experiment is indicated in the figure.

Statistical analysis was conducted using unpaired Student’s t test for two groups comparison (Figs 1C – 4C, D, G, H – 6C, D, G, H), one-way ANOVA followed up by Tukey’s multiple comparisons for three or more groups comparison (Figs. 1F – 2 – 7C, E), and two-way ANOVA followed up by Tukey’s multiple comparisons (Figs. 3B – 5 – 7A). All statistical comparisons were performed on the data from ≥ 3 independent samples. Significance is shown as *p < 0.05, **p < 0.01, ***p < 0.001, ****p < 0.0001.

## Supporting information

eLife author's response

## Acknowledgements

This work was funded in part by PHS NIH grants R01 AA027754 and PHS NIH R01 AG073900 (HY), as well as support the Professional Development Fund from the Geisel School of Medicine at Dartmouth (PM). We thank the Institute for Biomolecular Targeting at Dartmouth for technical assistance.

## Notes

Sources of support: This work was supported by PHS NIH R01 AA027754, PHS NIH R01 AG073900, and the Professional Development Fund granted by Geisel School of Medicine at Dartmouth. RK, MZ were supported in part by the Undergraduate Research Assistantships at Dartmouth College. MC was supported in part by the Dartmouth MD/PhD Undergraduate Summer Fellowship Program.

### Competing Interest Statement

The authors have declared no competing interest.

### Summary of Updates

New Figure 1 (experimental timeline and diet intake): We added a figure summarizing the experimental design, including maternal acclimation, PAE, and offspring assessments at each developmental stage. It also shows comparable maternal intake of ethanol-containing and control liquid diets, and clarifies the timing of the biochemical and behavioral assessments. Biochemical assessment at P0 (Figures 2 and 3): We revised the figure titles and text to reflect more accurately the effects of PAE on APP C-terminal fragments (APP-CTFs) and the Notch intracellular domain (NICD). The most robust PAE-associated changes occurred in wild-type mice, whereas differences between PAE-exposed and control 3xTg-AD mice did not reach statistical significance. Behavioral assessment at 3 months (Figure 4): We clarified the Morris water maze (MWM) results, emphasizing that learning impairment was primarily seen in PAE-exposed wild-type mice. We also explained that the non-exposed 3xTg-AD group serves as a reference for the behavioral effects of AD-associated genetic risk. Biochemical assessment at 3 months (Figure 5): We revised the text to emphasize the persistent increase in APP-CTFs after PAE in wild-type mice, and clarified the interpretation of the findings in the context of the 3xTg-AD genotype. Behavioral and biochemical assessments at 6 months (Figures 6 and 7): We refined the MWM interpretation to emphasize the pronounced spatial memory impairment in PAE-exposed wild-type mice. We also clarified the biochemical findings: a modest increase in hippocampal APP-CTFs after PAE, and more pronounced genotype-associated changes in 3xTg-AD mice. In addition, we added a direct statistical comparison between young and older PAE-exposed B6129 mice to evaluate the progression of behavioral deficits over time. PAE effects in 3xTg-AD mice (Figure 8): We revised the Results and Discussion to avoid overstating the impact of PAE on the 3xTg-AD behavioral phenotype. We now acknowledge that PAE did not significantly exacerbate learning deficits relative to non-exposed 3xTg-AD controls at four months. Although PAE-exposed 3xTg-AD mice showed a trend toward greater impairment, we have adjusted our conclusions to distinguish this trend from a statistically supported exacerbation of the genotype-associated phenotype. Data distribution by sex: We evaluated the distribution of data by sex to determine whether sex-related differences could influence the effects of PAE on γ-secretase-associated APP processing. Male and female animals were similarly distributed across experimental groups, and we found no apparent sex-related differences in PAE-associated changes in APP-CTFs.

